## Supplementary material for "Non-invasive Ultrasonic Neuromodulation of the Human Nucleus Accumbens Impacts Reward Sensitivity"

### Supplementary Results

#### TUS-dACC TUS and decision-making under low reward expectancy

The task design included trials where participants chose between two low-probability (30%) options, creating conditions of low expected value. These trials provide an opportunity to assess whether dACC stimulation modulates behaviour in contexts where adaptive choice requires evaluating similarly unrewarding alternatives. Previous studies in both humans[1,2] and macaques[3] suggest that the dACC is engaged in representing the value of counterfactual alternatives - options not chosen but potentially informative for future switching. Notably, in macaques, TUS-induced perturbation of dACC function abolishes the typical relationship between counterfactual value and behavioural change[3].

Building on this, we explored whether TUS-dACC might impair the brain's ability to recognize that the currently unchosen option is not meaningfully better than the chosen one. As a result, individuals might rely more heavily on broader indicators of low environmental reward—potentially mediated by dorsal raphe mechanisms [4,5] — and increase switching behaviour even when no advantageous alternative is available.

To test this, we conducted a targeted analysis of trials involving two low-probability stimuli. The results revealed that, after TUS-dACC, participants were more likely to switch choices following unrewarded outcomes in these trials compared to Sham. This finding supports the hypothesis that dACC stimulation disrupts the specific valuation of immediate alternatives, leading to greater reliance on global motivational signals that favour behavioural exploration. It further illustrates the nuanced role of dACC in value-guided learning and decision-making, particularly under conditions of ambiguity or uniformly low reward.

#### Supplementary Figure 1

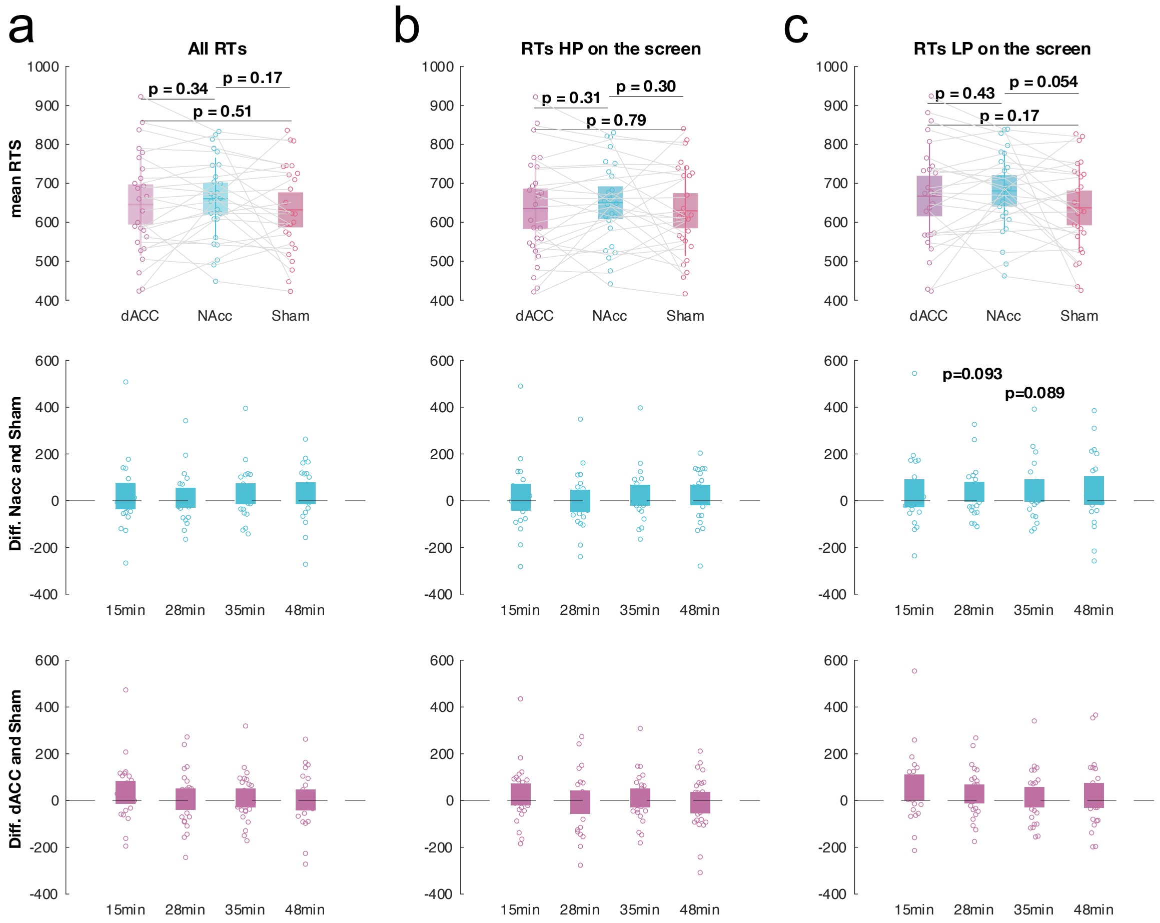


**Suppl. Fig. 1. Reaction time analyses across stimulation conditions and trial types. Top row:** Mean reaction times following TUS-dACC, TUS-NAcc, and Sham for all trials (left), HP trials (middle; trials with one high-probability and one low-probability option), and LP trials (right; trials with two low-probability options). Each dot represents an individual participant's mean across the task for the specified condition (n=26). Boxes represent the standard deviation around the mean. **Middle row:** Reaction times across the four post-TUS blocks (~15, 28, 35, and 48 minutes) as a difference between NAcc and Sham. Each dot represents an individual participant's mean (n=26). Boxes represent the standard deviation around the mean. **Bottom row:** Reaction times across the four post-TUS blocks (~15, 28, 35, and 48 minutes) and as a difference between dACC and Sham. Each dot represents an individual participant's mean (n=26). Boxes represent the standard deviation around the mean. For **a,b,c top row** statistical significance was determined using One-way ANOVA and two-sided t test. For **a,b,c middle and bottom row**, single two-sided t-tests were employed for each window. No multiple comparisons were applied.

#### Supplementary Figure 2

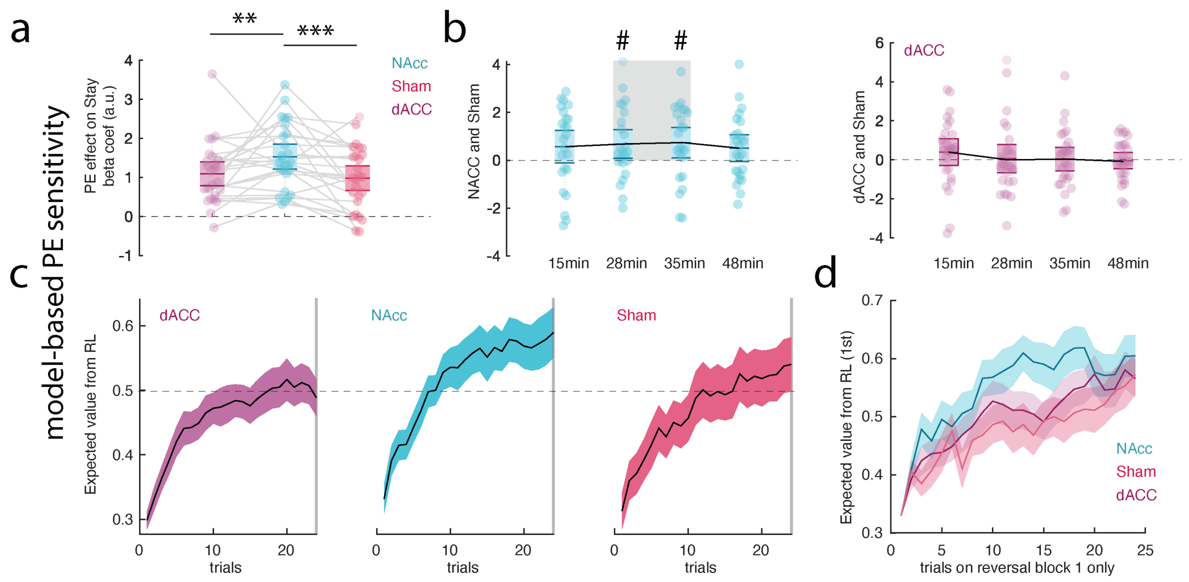


**Suppl. Fig. 2. Reinforcement learning model results. a** PE–stay analyses revealed an increase in the relationship between PE and subsequent stay behaviours after TUS-NAcc. Each dot represents an individual participant's mean beta estimate for the specified condition (n=26). **b** Left panel: Time course of TUS-NAcc effects on PE-related behaviour, showing the difference between TUS-NAcc and Sham conditions for each of the four post-TUS testing blocks. The TUS-NAcc-induced reward-related changes were most prominent in the middle of the approximately one-hour post-TUS period. This effect was not observed in the right panel, which shows the comparison between TUS-dACC and Sham; no significant time window emerged in that contrast (n=26). **c** Expected value from the RL models curves for the high-probability option across all reversal blocks. A 5-trial running average was applied to smooth trial-by-trial variability, which results in a slight shift in the apparent reversal point—appearing earlier than the actual reversal at trial 24 (trial 25 marks the start of the new reversal period). The shaded area represents the standard error of the mean. **d** Same as **c** but showing only the first reversal block. The three conditions (TUS-NAcc, TUS-dACC, and Sham) are presented stacked and with transparency to allow for direct visual comparison. Exact p values are presented in the supplementary table 5. **a**: statistical significance was determined using One-way ANOVA and two-sided t tests. **b**: single two-sided t-tests were employed for each window. No multiple comparisons were applied.

#### Supplementary Figure 3

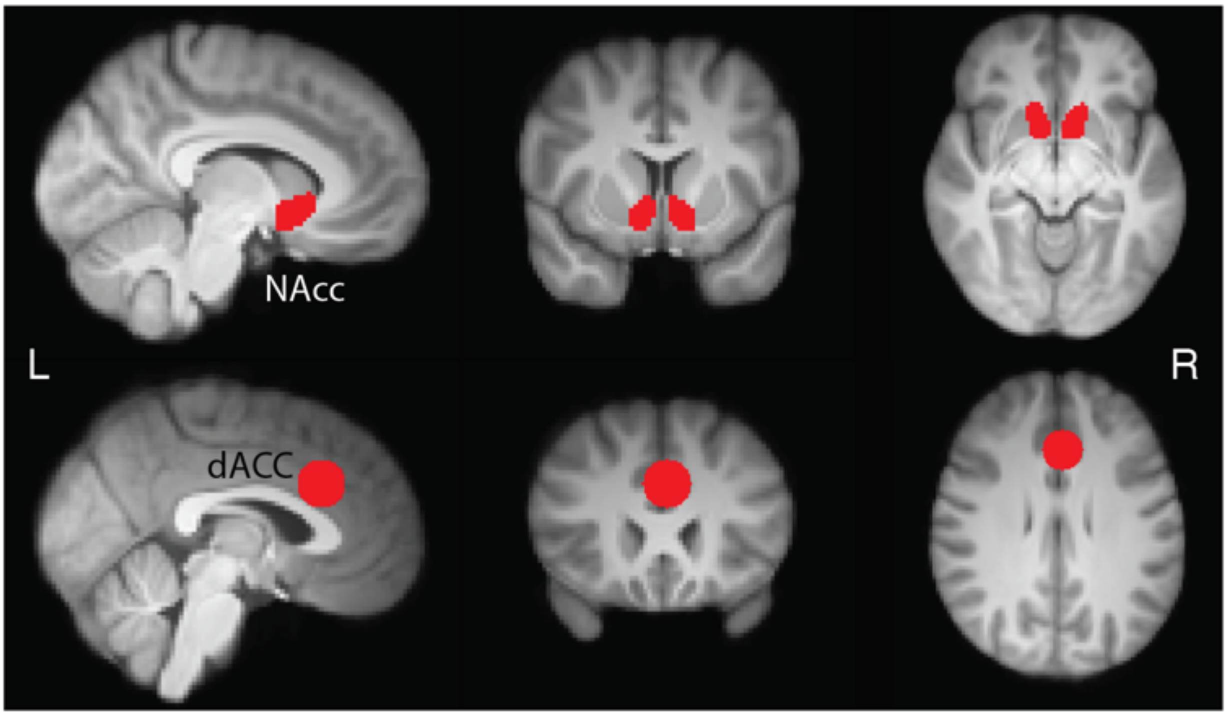


**Suppl. Fig. 3.** Region of interest (ROI) definition for the ROI analysis reported in the FMRI findings section for both the NAcc and the dACC regions. ROI are presented in red.

### Supplementary Figure 4

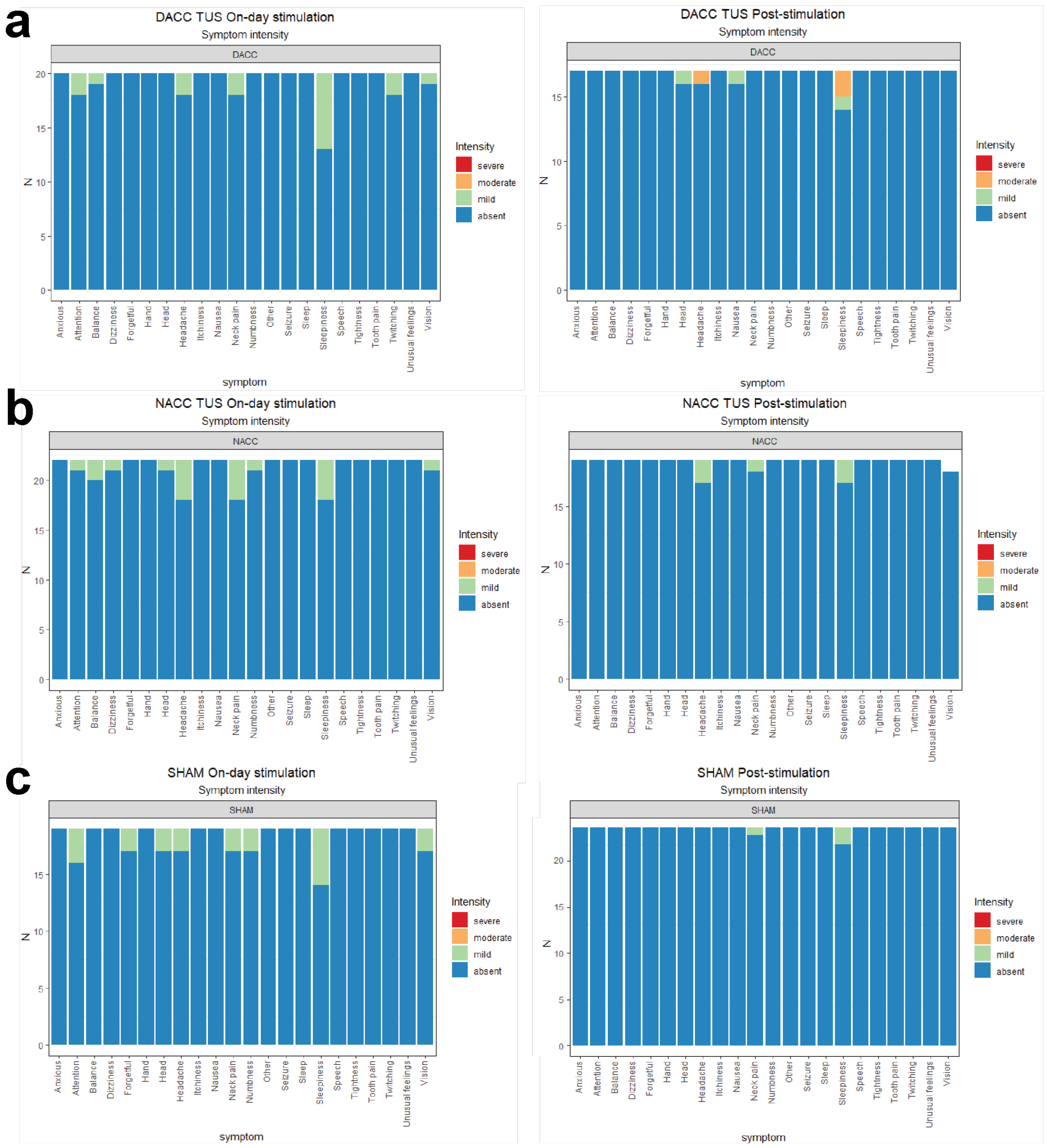


**Suppl. Fig. 4. Group safety report. a** Total number of responses for all participants (n = 26) from the dACC condition on the day (left) and on the following day (right), coded by the severity of the symptom. **b** Total number of responses for all participants (n = 26) from the NAcc condition on the day (left) and on the following day (right), coded by the severity of the symptom. **c** Total number of responses for all participants (n = 26) from the Sham condition on the day (left) and on the following day (right), coded by the severity of the symptom. Note: Unusual feelings refer to the question asking about experiencing any unusual feelings, attitudes or emotions.

#### Supplementary Table 1 (NAcc)

**Suppl. Table 1.** Acoustic simulation parameters and output for all study participants in the Nacc condition. The table reports the pressure values at the spatial-peak (Max Pressure), corresponding transcranial mechanical index (MItc) and spatial-peak pulse-average intensity (ISPPA), along with the maximum temperature rise in the head, to assess safety. To evaluate stimulation efficacy, the in situ estimate for the pressure amplitude at the target, Isppa at the target and the size of the -6dB focal volume are reported. Pressure values are given in megapascals (MPa)

| Focus depth | Max Pressure (MPa) | MItc | Isppa (W/cm2) | Max. temp. rise | Pressure at target (MPa) | Isppa at target (W/cm2) | -6dB focal volume (mm3) |
|---|---|---|---|---|---|---|---|
| 82 | 0.44 | 0.63 | 6.54 | 3.37 | 0.32 | 3.43 | 561.13 |
| 71 | 0.64 | 0.90 | 13.46 | 2.14 | 0.43 | 6.06 | 257.75 |
| 74 | 0.66 | 0.94 | 14.62 | 2.31 | 0.30 | 2.98 | 234.75 |
| 74 | 0.66 | 0.93 | 14.41 | 2.70 | 0.46 | 6.99 | 283.63 |
| 82 | 0.62 | 0.88 | 12.79 | 2.39 | 0.56 | 10.56 | 458.13 |
| 82 | 0.61 | 0.87 | 12.53 | 4.59 | 0.48 | 7.61 | 327.00 |
| 75 | 0.62 | 0.87 | 12.76 | 3.07 | 0.51 | 8.71 | 281.75 |
| 77 | 0.61 | 0.86 | 12.25 | 3.61 | 0.55 | 10.18 | 332.00 |
| 80 | 0.63 | 0.89 | 13.17 | 3.05 | 0.46 | 7.04 | 303.63 |
| 74 | 0.65 | 0.92 | 14.21 | 2.83 | 0.19 | 1.23 | 216.00 |
| 76 | 0.64 | 0.90 | 13.64 | 3.10 | 0.55 | 10.00 | 255.50 |
| 77 | 0.52 | 0.74 | 9.11 | 3.13 | 0.40 | 5.31 | 490.25 |
| 69 | 0.66 | 0.94 | 14.61 | 1.34 | 0.53 | 9.42 | 178.50 |
| 74 | 0.53 | 0.75 | 9.25 | 2.46 | 0.35 | 4.20 | 406.25 |
| 74 | 0.59 | 0.84 | 11.76 | 2.11 | 0.29 | 2.79 | 367.63 |
| 73 | 0.63 | 0.90 | 13.42 | 2.59 | 0.54 | 9.89 | 282.63 |
| 72 | 0.60 | 0.84 | 11.82 | 2.19 | 0.50 | 8.29 | 329.00 |
| 74 | 0.59 | 0.83 | 11.51 | 1.35 | 0.45 | 6.81 | 379.38 |
| 80 | 0.56 | 0.80 | 10.56 | 1.89 | 0.25 | 2.04 | 450.38 |
| 82 | 0.63 | 0.89 | 13.07 | 3.88 | 0.56 | 10.64 | 420.38 |
| 74 | 0.64 | 0.91 | 13.86 | 1.71 | 0.37 | 4.52 | 200.38 |
| 79 | 0.67 | 0.94 | 14.82 | 3.27 | 0.63 | 13.41 | 351.75 |
| 76 | 0.57 | 0.80 | 10.70 | 3.04 | 0.52 | 9.18 | 393.25 |
| 73 | 0.64 | 0.91 | 13.81 | 1.71 | 0.32 | 3.46 | 211.00 |
| 79 | 0.61 | 0.86 | 12.44 | 2.50 | 0.41 | 5.53 | 358.63 |
| 75 | 0.68 | 0.95 | 15.19 | 1.82 | 0.53 | 9.19 | 197.25 |

#### Supplementary Table 2 (dACC)

**Suppl. Table 2.** Acoustic simulation parameters and output for all study participants in the dACC condition. The table reports the pressure values at the spatial-peak (Max Pressure), corresponding transcranial mechanical index (MItc) and spatial-peak pulse-average intensity (ISPPA), along with the maximum temperature rise in the head, to assess safety. To evaluate stimulation efficacy, the in situ estimate for the pressure amplitude at the target, Isppa at the target and the size of the -6dB focal volume are reported. Pressure values are given in megapascals (MPa)

| Focus depth | Max Pressure (MPa) | MItc | Isppa (W/cm2) | Max. temp. rise | Pressure at target (MPa) | Isppa at target (W/cm2) | -6dB focal volume (mm3) |
|---|---|---|---|---|---|---|---|
| 64.00 | 0.64 | 0.90 | 13.59 | 1.95 | 0.59 | 11.60 | 175.75 |
| 57.00 | 0.64 | 0.90 | 13.61 | 0.93 | 0.52 | 9.01 | 113.63 |
| 57.00 | 0.52 | 0.73 | 8.86 | 0.94 | 0.41 | 5.60 | 210.38 |
| 61.00 | 0.61 | 0.86 | 12.31 | 1.09 | 0.49 | 8.00 | 175.13 |
| 57.00 | 0.65 | 0.92 | 14.06 | 0.85 | 0.52 | 9.01 | 132.38 |
| 58.00 | 0.57 | 0.81 | 10.99 | 1.01 | 0.39 | 5.07 | 166.13 |
| 57.00 | 0.57 | 0.80 | 10.78 | 0.90 | 0.37 | 4.56 | 167.63 |
| 54.00 | 0.61 | 0.86 | 12.28 | 0.73 | 0.60 | 12.00 | 115.13 |
| 58.00 | 0.47 | 0.66 | 7.21 | 1.21 | 0.43 | 6.16 | 278.38 |
| 57.00 | 0.51 | 0.72 | 8.67 | 1.01 | 0.37 | 4.56 | 201.88 |
| 61.00 | 0.55 | 0.78 | 10.26 | 1.29 | 0.42 | 5.88 | 207.50 |
| 58.00 | 0.53 | 0.75 | 9.41 | 0.88 | 0.41 | 5.60 | 207.00 |
| 57.00 | 0.62 | 0.87 | 12.74 | 0.81 | 0.53 | 9.36 | 131.88 |
| 61.00 | 0.65 | 0.92 | 14.03 | 1.28 | 0.52 | 9.01 | 144.00 |
| 61.00 | 0.64 | 0.90 | 13.47 | 1.39 | 0.43 | 6.16 | 158.75 |
| 60.00 | 0.62 | 0.88 | 12.98 | 1.16 | 0.51 | 8.67 | 165.13 |
| 53.00 | 0.55 | 0.78 | 10.09 | 0.95 | 0.43 | 6.16 | 163.63 |
| 59.00 | 0.61 | 0.86 | 12.42 | 1.00 | 0.40 | 5.33 | 157.25 |
| 62.00 | 0.45 | 0.63 | 6.67 | 1.16 | 0.35 | 4.08 | 92.63 |
| 62.00 | 0.64 | 0.90 | 13.54 | 1.35 | 0.47 | 7.36 | 159.38 |
| 65.00 | 0.49 | 0.69 | 7.97 | 1.63 | 0.39 | 5.07 | 128.25 |
| 69.00 | 0.61 | 0.87 | 12.50 | 1.46 | 0.48 | 7.68 | 84.25 |
| 63.00 | 0.56 | 0.80 | 10.58 | 1.20 | 0.54 | 9.72 | 196.13 |
| 60.00 | 0.65 | 0.93 | 14.29 | 1.09 | 0.56 | 10.45 | 141.63 |
| 58.00 | 0.54 | 0.76 | 9.57 | 1.21 | 0.38 | 4.81 | 196.13 |
| 64.00 | 0.52 | 0.73 | 8.98 | 1.57 | 0.48 | 7.68 | 262.75 |

#### Supplementary Table 3

**Suppl. Table 3.** ANOVA results for no reinforcement learning model: effect of reward on Win–Stay. Post-hoc t-tests revealed a stronger relationship between reward and subsequent win–stay behaviour after TUS-NAcc compared to Sham. No differences were observed for TUS-dACC compared to Sham. The results for each task block are also shown.

**ANOVA**

| Sum of Squares | DF | Mean Squares | F | p-value |
|---|---|---|---|---|
| 6.4468 | 2 | 3.2234 | 3.2954 | **0.0424** |
| 73.3613 | 75 | 0.9781 | | |
| 79.8082 | 77 | | | |

**Post hoc t-tests**

| | p-value | CI low | CI high | t-stat | DF | SD |
|---|---|---|---|---|---|---|
| NAcc-Sham | **0.0110** | -0.9550 | -0.1364 | -2.7461 | 25 | 1.0133 |
| NAcc-dACC | **0.0016** | -1.0435 | -0.2730 | -3.5193 | 25 | 0.9537 |
| dACC-Sham | 0.5732 | -0.2935 | 0.5186 | 0.5707 | 25 | 1.0054 |

**Difference NAcc and Sham**

| | p-value | CI low | CI high | t-stat | DF | SD |
|---|---|---|---|---|---|---|
| 15min | 0.0867 | -0.0936 | 1.2942 | 1.7894 | 23 | 1.6434 |
| 28min | **0.0056** | 0.3496 | 1.8249 | 3.0490 | 23 | 1.7469 |
| 35min | 0.0537 | -0.0119 | 1.3766 | 2.0330 | 23 | 1.6442 |
| 48min | 0.1133 | -0.1453 | 1.2784 | 1.6462 | 23 | 1.6859 |

**Difference dACC and Sham**

| | p-value | CI low | CI high | t-stats | DF | SD |
|---|---|---|---|---|---|---|
| 15min | 0.4118 | -0.4827 | 1.1404 | 0.8345 | 25 | 2.0093 |
| 28min | 0.6233 | -0.5226 | 0.8553 | 0.4972 | 25 | 1.7058 |
| 35min | 0.7145 | -0.6919 | 0.9949 | 0.3699 | 25 | 2.0881 |
| 48min | 0.4832 | -0.7650 | 0.3720 | -0.7117 | 25 | 1.40764 |

### Supplementary Table 4

**Suppl. Table 4.** Comparison of learning curves across trials and reversal periods. Participants were more likely to select the high probability option at the end of a reversal period after TUS-NAcc compared to TUS-dACC and Sham. This is particularly true for the first reversal of the block.

**All reversal periods**

| Fixed Effect | Estimate | SE | t-value | DF | p-value |
|---|---|---|---|---|---|
| trial | 0.0199 | 0.0026 | 7.6579 | 5924 | 2.19E-14 |
| cond | -0.0113 | 0.0041 | -2.7408 | 5924 | **0.0061** |
| trial^2 | -0.0006 | 0.0001 | -5.0510 | 5924 | 4.53E-07 |

**First reversal period only**

| Fixed Effect | Estimate | SE | t-value | DF | p-value |
|---|---|---|---|---|---|
| trial | 0.0226 | 0.0033 | 6.8422 | 1478 | 1.14E-11 |
| cond | -0.0208 | 0.0052 | -3.9714 | 1478 | **7.49E-05** |
| trial^2 | -0.0008 | 0.0001 | -5.3668 | 1478 | 9.29E-08 |

#### Supplementary Table 5

**Suppl. Table 5.** ANOVA results for reinforcement learning model: effect of prediction error (PE) on Win–Stay. Post-hoc t-tests revealed a stronger relationship between PE and subsequent win–stay behaviour after TUS-NAcc compared to Sham. No differences were observed for TUS-dACC compared to Sham. The results for each task block are also shown.

**ANOVA**

| Sum of Squares | DF | Mean Squares | F | p-value |
|---|---|---|---|---|
| 4.4277 | 2 | 2.2138 | 3.3304 | **0.0411** |
| 49.8554 | 75 | 0.6647 | | |
| 54.2831 | 77 | | | |

**Post hoc t-tests**

| | p-value | CI low | CI high | t-stat | DF | SD |
|---|---|---|---|---|---|---|
| NAcc-Sham | **0.0274** | -0.8249 | -0.0532 | -2.3433 | 25 | 0.9554 |
| NAcc-dACC | **0.0031** | -0.8996 | -0.2054 | -3.2786 | 25 | 0.8593 |
| dACC-Sham | 0.5138 | -0.2393 | 0.4662 | 0.6624 | 25 | 0.8733 |

**Difference NAcc and Sham**

| | p-value | CI low | CI high | t-stat | DF | SD |
|---|---|---|---|---|---|---|
| 15min | 0.1142 | -0.1476 | 1.2837 | 1.6421 | 23 | 1.6948 |
| 28min | **0.0323** | 0.0680 | 1.4064 | 2.2788 | 23 | 1.5849 |
| 35min | **0.0360** | 0.0482 | 1.3102 | 2.2267 | 23 | 1.4943 |
| 48min | 0.0865 | -0.0792 | 1.0995 | 1.7908 | 23 | 1.3957 |

**Difference dACC and Sham**

| | p-value | CI low | CI high | t-stat | DF | SD |
|---|---|---|---|---|---|---|
| 15min | 0.2647 | -0.3215 | 1.1201 | 1.1408 | 25 | 1.7847 |
| 28min | 0.9063 | -0.5977 | 0.6709 | 0.1189 | 25 | 1.5705 |
| 35min | 0.8685 | -0.7021 | 0.8262 | 0.1673 | 25 | 1.8919 |
| 48min | 0.8369 | -0.4817 | 0.3933 | -0.2080 | 25 | 1.0831 |

### Supplementary Table 6

**Suppl. Table 6.** Comparison of learning curves (that is the rate of choosing the high probability option over the period of one reversal) between TUS-NAcc, TUS-dACC, and Sham.

**All reversal periods**

| Fixed Effect | Estimate | SE | t-value | DF | p-value |
|---|---|---|---|---|---|
| trial | 0.0300 | 0.0014 | 21.9431 | 7785 | **1E-103** |
| cond | 0.0056 | 0.0029 | 1.9352 | 7785 | 0.0529 |
| trial^2 | -0.0009 | 0.0001 | -18.3870 | 7785 | **6E-74** |

**First reversal period only**

| Fixed Effect | Estimate | SE | t-value | DF | p-value |
|---|---|---|---|---|---|
| trial | 0.0192 | 0.0021 | 9.3135 | 1867 | **3E-20** |
| cond | -0.0117 | 0.0042 | -2.7863 | 1867 | **0.0053** |
| trial^2 | -0.0005 | 0.0001 | -5.7059 | 1867 | **1E-08** |

#### Supplementary Table 7

**Suppl. Table 7.** ANOVA and post-hoc test results for learning rates linked with positive and negative feedback. The learning rates linked with positive feedback were higher after TUS-NAcc compared to dACC. The learning rates associated with negative feedback were not significantly different across the conditions.

**ANOVA**

**Learning rate (positive feedback)**

| Sum of Squares | DF | Mean Squares | F | p-value |
|---|---|---|---|---|
| 0.3760 | 2 | 0.18802 | 5.6847 | **0.0050** |
| 2.4806 | 75 | 0.03307 | | |
| 2.8567 | 77 | | | |

**Learning rate (negative feedback)**

| Sum of Squares | DF | Mean Squares | F | p-value |
|---|---|---|---|---|
| 0.0246 | 2 | 0.0123 | 0.2758 | 0.7597 |
| 3.3508 | 75 | 0.0446 | | |
| 3.3754 | 77 | | | |

**Post-hoc tests for learning rate (positive feedback)**

| | p-value | CI low | CI high | t-stat | DF | SD |
|---|---|---|---|---|---|---|
| NAcc-Sham | **0.0211** | -0.2182 | -0.0194 | -2.4605 | 25.0000 | 0.2462 |
| NAcc-dACC | **0.0060** | -0.2778 | -0.0518 | -3.0031 | 25.0000 | 0.2798 |
| dACC-Sham | 0.3432 | -0.0521 | 0.1441 | 0.9662 | 25.0000 | 0.2429 |

#### Supplementary Table 8

**Suppl. Table 8.** ANOVA results for win–stay strategy and maladaptive choices. We found a main effect of condition on the rate of maladaptive choices after TUS-NAcc.

**ANOVA**
**Win–stay after low probability outcome**

| Sum of Squares | DF | Mean Squares | F | p-value |
|---|---|---|---|---|
| 8.4220 | 2 | 4.2110 | 3.7525 | **0.0279** |
| 84.1643 | 75 | 1.1221 | | |
| 92.5864 | 77 | | | |

**Single t-test against 0**

| | p-value | CI low | CI high | t-stat | DF | SD |
|---|---|---|---|---|---|---|
| dACC | 0.3988 | -0.2630 | 0.6390 | 0.8584 | 25 | 1.1166 |
| NAcc | **0.0033** | 0.2675 | 1.1978 | 3.2442 | 25 | 1.1515 |
| Sham | 0.7646 | -0.4126 | 0.3069 | -0.3026 | 25 | 0.89079 |

### Supplementary Table 9

**Suppl. Table 9.** ROI analysis results for PE during reward delivery and reward expectation. There was an increase in the parametric response to reward expectation in the NAcc in the TUS-NAcc condition compared to the Sham and TUS-dACC conditions.

**ANOVA**

**NAcc reward expectation**

| Sum of Squares | DF | Mean Squares | F | p-value |
|---|---|---|---|---|
| 2.84814906 | 2 | 1.4240 | 7.1545 | **0.0014** |
| 14.9283081 | 75 | 0.1990 | | |
| 17.7764571 | 77 | | | |

**dACC reward delivery**

| Sum of Squares | DF | Mean Squares | F | p-value |
|---|---|---|---|---|
| 1.6866 | 2 | 0.8433 | 3.1237 | **0.0497** |
| 20.2483 | 75 | 0.2699 | | |
| 21.9349 | 77 | | | |

**Post hoc t-tests NAcc reward expectation**

| | p-value | CI low | CI high | t-stat | DF | SD |
|---|---|---|---|---|---|---|
| dACC-NAcc | **0.0006** | -0.7057 | -0.2193 | -3.9165 | 25 | 0.6021 |
| NAcc-Sham | **0.0248** | -0.5469 | -0.0405 | -2.3888 | 25 | 0.6269 |
| dACC-Sham | 0.1918 | -0.4280 | 0.0903 | -1.3416 | 25 | 0.6416 |

**Post hoc t-tests dACC reward delivery**

| | p-value | CI low | CI high | t-stat | DF | SD |
|---|---|---|---|---|---|---|
| dACC-NAcc | **0.0076** | -0.6140 | -0.1042 | -2.9023 | 25 | 0.6309 |
| NAcc-Sham | 0.2017 | -0.4004 | 0.0889 | -1.3110 | 25 | 0.6058 |
| dACC-Sham | 0.1514 | -0.4864 | 0.0796 | -1.4797 | 25 | 0.7008 |

### Supplementary Table 10

**Suppl. Table 10.** Demographic information about the three DBS patients. DBS: deep brain stimulation; OCD: obsessive–compulsive disorder; MDD: major depressive disorder.

| Patient | Sex | Age | Psychiatric comorbidities | Bilateral DBS Settings | | |
|---|---|---|---|---|---|---|
| | | | | Voltage (V) | Pulse Width (μs) | Frequency (Hz) |
| 1 | F | 31 | OCD | 3.5 | 90 | 130 |
| 2 | F | 34 | OCD<br>MDD | 4.0 | 80 | 130 |
| 3 | F | 61 | OCD<br>MDD | 4.0 | 60 | 130 |
| Mean (SD) | | 42 (13.5) | | | | |

### Supplementary Material 1

#### *Resource Table*

| RESOURCE | SOURCE | IDENTIFIER |
| --- | --- | --- |
| **Deposited data** | | |
| Structural and functional MRI data ("TUS fMRI NAcc part 1-3")<br>Behavioural data | Siti Yaakub<br>Elsa Fouragnan | https://osf.io/j34qz/<br>https://osf.io/w3mev/<br>https://osf.io/vst9y/ |
| **Software and algorithms** | | |
| MATLAB R2023a | Mathworks | https://www.mathworks.com/ |
| FMRIB Software Library v6.0 | FMRIB, Oxford, UK | https://fsl.fmrib.ox.ac.uk/ |
| HD-BET | Github | https://github.com/MIC-DKFZ/HD-BET |
| k-Wave toolbox v1.4 | Github | http://www.k-wave.org |
| MR-to-pCT toolbox v1 | Github | https://github.com/sitiny/mr-to-pct |
| BRIC TUS Simulation Tools v2 | Github | https://github.com/sitiny/BRIC_TUS_Simulation_Tools |
| Reinforcement learning | Github | https://github.com/efouragnan/RL_models |
| Presentation Neurobs | Presentation | https://www.neurobs.com/ |
| **Other** | | |
| NeuroFUS CTX-500 with optimized steering range (40–80mm) | BrainBox, SonicConcepts | https://brainbox-neuro.com/products/neurofus<br>http://sonicconcepts.com |
| Stereotaxic neuronavigation system | Brainsight | https://www.rogue-research.com/tms/brainsight-tms/ |
| Siemens Prima (MR) | BRIC MRI | https://www.plymouth.ac.uk/facilities/brain-research-imaging-centre<br>https://www.siemens-healthineers.com/magnetic-resonance-imaging/3t-mri-scanner/magnetom-prisma |

### Supplementary Material 2

#### *Safety Questionnaire*

Please rate the intensity of any symptoms you have had since your stimulation according to the following criteria:
**Absent** = Not present
**Mild** = Present but not bothersome
**Moderate** = Tolerable—required some intervention/medication, but did not interfere with day-to-day activities
**Severe** = Intolerable—required contact with a GP or hospital A&E

Use the box to the right of each symptom to provide details, e.g., describe what you felt, how long it lasted, medication you took to relieve the symptoms, and whether you think it might be related to the stimulation.

| **Since your stimulation, have you had ... ?** | **Intensity of symptom** | **Please provide details** |
|---|---|---|
| a headache | ☐ absent ☐ moderate ☐ mild ☐ severe | |
| neck pain | ☐ absent ☐ moderate ☐ mild ☐ severe | |
| tooth pain | ☐ absent ☐ moderate ☐ mild ☐ severe | |
| unusual feelings on your head or scalp | ☐ absent ☐ moderate ☐ mild ☐ severe | |
| itchiness | ☐ absent ☐ moderate ☐ mild ☐ severe | |
| changes to your hearing | ☐ absent ☐ moderate ☐ mild ☐ severe | |
| speech problems | ☐ absent ☐ moderate ☐ mild ☐ severe | |
| vision problems (e.g., double vision) | ☐ absent ☐ moderate ☐ mild ☐ severe | |
| unusual twitching or muscle movement | ☐ absent ☐ moderate ☐ mild ☐ severe | |
| difficulties in balance | ☐ absent ☐ moderate ☐ mild ☐ severe | |
| changes in the movement of your strongest hand | ☐ absent ☐ moderate ☐ mild ☐ severe | |
| numbness or tingling sensations | ☐ absent ☐ moderate ☐ mild ☐ severe | |
| muscle tightness of the face or arm | ☐ absent ☐ moderate ☐ mild ☐ severe | |
| unusual feelings, attitudes or emotions | ☐ absent ☐ moderate ☐ mild ☐ severe | |
| anxiety, worried thoughts or nervousness | ☐ absent ☐ moderate ☐ mild ☐ severe | |
| increased sleepiness | ☐ absent ☐ moderate ☐ mild ☐ severe | |
| changes to your sleep pattern | ☐ absent ☐ moderate ☐ mild ☐ severe | |
| difficulty paying attention | ☐ absent ☐ moderate ☐ mild ☐ severe | |
| increased forgetfulness | ☐ absent ☐ moderate ☐ mild ☐ severe | |
| nausea or sickness to the stomach | ☐ absent ☐ moderate ☐ mild ☐ severe | |
| dizziness or light-headedness | ☐ absent ☐ moderate ☐ mild ☐ severe | |
| a seizure within the last 24 hours | ☐ absent ☐ moderate ☐ mild ☐ severe | |
| other symptoms: ____________________ | ☐ absent ☐ moderate ☐ mild ☐ severe | |

Do you have anything else to report? ________________________
